# Evolutionarily conserved midbody reorganization precedes ring canal formation during gametogenesis

**DOI:** 10.1101/2022.06.03.494691

**Authors:** Kari L. Price, Dyuthi M. Tharakan, Lynn Cooley

## Abstract

How canonical cytokinesis is altered during germ cell division to produce stable intercellular bridges called ring canals is poorly understood. Here, using time-lapse imaging in *Drosophila*, we observe that ring canal formation occurs via reorganization of the germline mid-body, a structure classically associated with its function in recruiting abscission-regulating proteins in complete cytokinesis. Germline midbodies reorganize from a focus into a ring rather than being discarded, and this transition is accompanied by changes in centralspindlin dynamics. The midbody-to-ring canal transformation is conserved in the *Drosophila* male and female germlines and during mouse and *Hydra* spermatogenesis. In *Drosophila*, ring canal formation depends on Citron kinase function in stark contrast to its role in abscission during somatic cell cytokinesis. Our results challenge existing models of ring canal formation, and provide important insights into broader functions of incomplete cytokinesis events across biological systems, such as those observed during development and disease states.

---

**I**n contrast to canonical cell division, animal germline cells and certain somatic cells undergo incomplete cytokinesis to produce cells in syncytial groups that remain connected by intercellular bridges, often called ring canals. Although intercellular bridges are conserved in male and female germline cells from sponges to mammals (1–8), we know relatively little about how cytokinesis is altered during germ cell division to give rise to these bridges.

Complete cytokinesis is orchestrated by a specific organelle, the midbody (9). The midbody resides at the center of the transient intercellular bridge between nascent cells and serves to recruit microtubule severing proteins and the membrane abscission machinery needed to finally separate new sister cells (10–12). Contrary to this function, certain studies implicate a possible relationship between germline midbodies and incomplete cell division resulting in ring canals. By electron microscopy, ring canals are characterized by an electron dense limiting membrane that resembles the cell furrow at the midbody stage of cytokinesis (13, 14). In the mouse male germline, loss of the ring canal-promoting factor TEX14, which blocks abscission through competitive inhibition of the ESCRT-associated protein CEP-55 (15), results in the formation of midbody-like foci between dividing calls, implying that midbody formation is a normal step in ring canal biogenesis (16). However, a CEP-55-based mechanism to block abscission is restricted to mammals since more primitive animal genomes do not encode CEP-55. A different model has emerged from studies of fixed *Drosophila* cells in which ring canals form due to arrested constriction of the contractile ring before midbody formation (5, 6, 8, 17–19).

The temporal aspects of ring canal formation have not been described and it remains unclear how midbody or contractile ring behavior during incomplete cytokinesis contributes to the formation of a stable ring canal. Furthermore, compositional differences between *Drosophila* male and female germline ring canals and between mouse and *Drosophila* male ring canals that, combined with differences in proposed models of ring canal formation, raise the question of whether there is one conserved mechanism of ring canal formation.

To gain new insight into the mechanism of ring canal formation, we performed live imaging to monitor the dynamics of ring canal proteins during incomplete mitosis and subsequent ring canal formation. We imaged several cleavage furrow and ring canal proteins, including the highly conserved MKLP1/kinesin-6 subunit of the centralspindlin complex. In both the *Drosophila* testis and the ovary, we discover that ring canal formation is preceded by the formation of a mid-body intermediate that, rather than being discarded, reorganizes to form a ring with open lumen. Further, by analysing MKLP1/kinesin-6 homologs in the mouse and *Hydra vulgaris* testis, we find that a midbody-to-ring canal transition is a conserved feature of ring canal formation across evolution. Mechanistically, we demonstrate a key role for the conserved serine-threonine kinase Citron kinase, called Sticky in *Drosophila*, in the midbody-to-ring canal transition. Our results reveal a previously unknown, but highly conserved midbody behavior during programmed incomplete cytokinesis and new insights into the mechanisms that distinguish complete versus incomplete cytokinesis.

## Results

### *Drosophila* ring canal formation during gametogenesis occurs via a midbody-like intermediate

To investigate how germline ring canals are formed, we performed time-lapse confocal imaging of known ring canal components during incomplete mitosis in the *Drosophila* germline (Fig. 1A)(Extended Data Fig. 1). We focused on the highly conserved MKLP1/kinesin-6 protein (known as Pavarotti or Pav in *Drosophila* and MKLP1/KIF23 in mammals), a subunit of the centralspindlin complex and component of all characterized ring canals (5, 6). In *Drosophila*, Pav has also been shown to localize to the central spindle and nascent cleavage furrow (20). We imaged either endogenously-tagged Pav::mCherry (see Materials and Methods) or a Pav::GFP transgenic protein expressed at endogenous levels (21). To correlate ring canal formation with cell cycle timing, we visualized Pav::mCherry in combination with GFP::Histone. Live imaging revealed that, during anaphase, Pav::mCherry localized to the cleavage furrow and central spindle as expected, consistent with its role in cytokinetic furrowing (Fig. 1B, t=0 min and 1C, t=0 min). However, rather than a cleavage furrow arrest as predicted by the current model for ring canal formation in *Drosophila* (6), we observed full constriction of the contractile ring and formation of a dense midbody-like focus containing Pav::mCherry (Fig. 1B, t=4 min). The midbody-like focus persisted for 25 minutes before resolving into a ring with an open lumen (Fig. 1B, t=34 minutes). During the period from nascent midbody to ring canal, the fluorescence intensity of Pav diminished, suggesting a period of midbody maturation (Fig. 1C). Pav pixel intensity during incomplete cytokinesis was highest in the nascent midbody focus and underwent a 2-fold reduction in fluorescence intensity over a period of 10 minutes (Fig. 1D). Resolution to a nascent ring canal with an open lumen was accompanied by an additional 3-fold reduction in fluorescence. We found that midbody formation accompanied every incomplete mitotic event in the testis, from gonialblast to 8-cell cyst divisions. Interestingly, the centralspindlin foci formed during incomplete cytokinesis were larger in diameter than the presumed midbody remnants formed following complete cytokinesis in the testis hub (compare 1C with 1E) revealing inherent differences in midbody morphology between cells that do or do not undergo abscission.

**Fig. 1.**
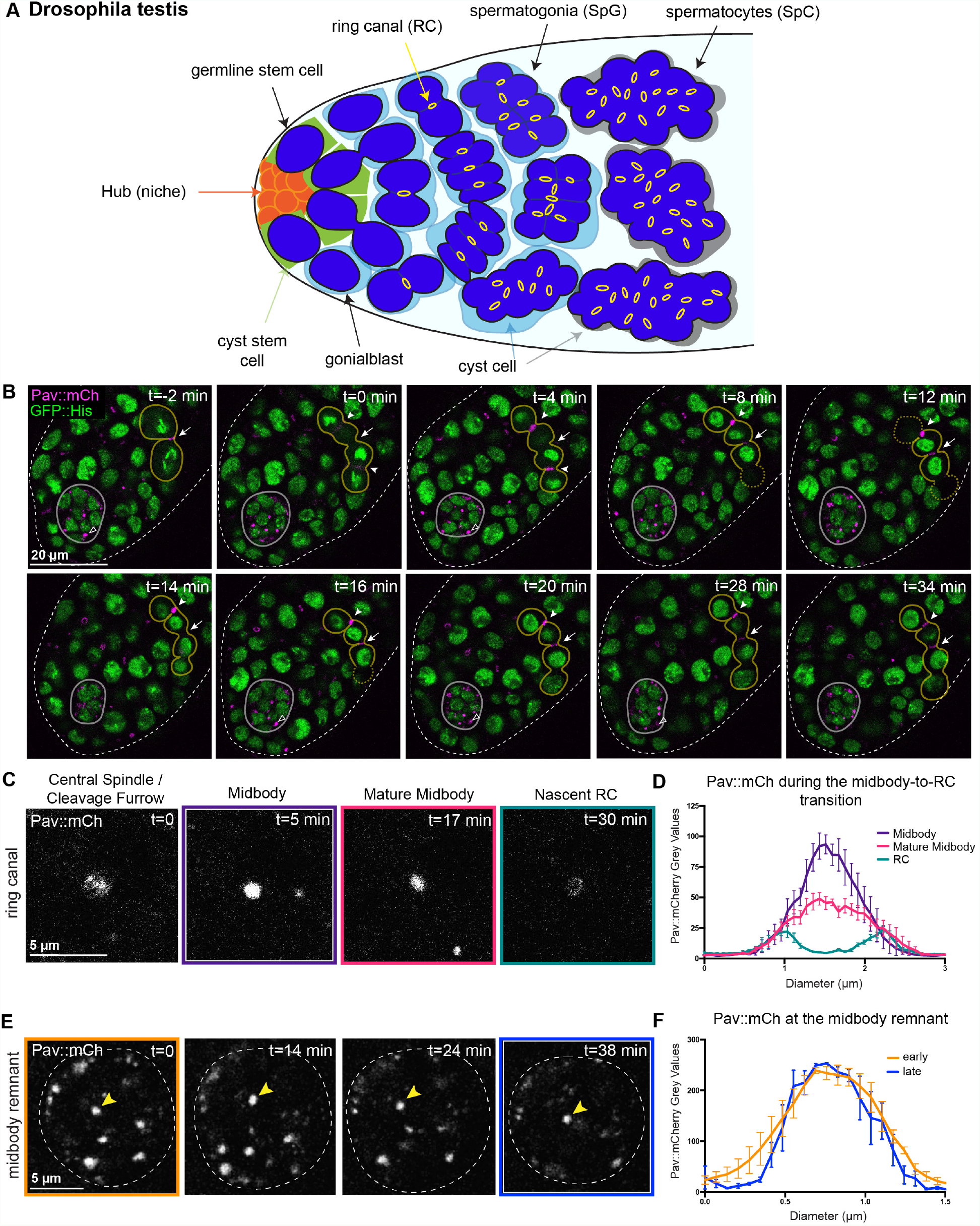
Ring canal formation in the *Drosophila* testis occurs via a midbody-like intermediate. (A) Schematic of the apical tip of the Drosophila testis. (B) Single Z-slices from a time-lapse movie of a *Drosophila* testis expressing Pavarotti (Pav)::mCh and GFP::Histone. The hub/stem cell niche (outlined in grey) is located at the apical tip of the testis; an open arrowhead marks a Pav-labeled midbody remnant (MR). An actively dividing two-cell germline cyst (outlined in yellow) is visible with a single ring canal (white arrow). (t=0 minutes) Pav::mCh is present at the cleavage furrow and central spindle (white arrowheads) of newly dividing cells. (t=4 minutes) Pav::mCh marks the constricting contractile ring. (t=8 minutes) Following constriction, Pav::mCh appears as a midbody (only one midbody is in focus; the dotted line marks the boundaries of the cell that are out of the plane of focus). The midbody focus persists from t=8 minutes to t=20 minutes, following which it begins to elongate (t=28 minutes) before appearing as a ring in the transverse orientation (t=34 minutes). (C) The stages of ring canal formation as revealed by Pav::mCh. (D) Pav::mCh pixel intensity values from line traces across the midbody and ring canal at each stage of ring canal formation (n=6; error bars display SEM). (E) Pav::mCh in midbody remnants in the hub. (F) Pav::mCh fluorescence intensity in midbody remnants (n=4).

Based on our live imaging of Pav::mCherry, we defined four stages of ring canal biogenesis as: 1) central spindle formation/cleavage furrow ingression; 2) midbody formation; 3) midbody maturation; and 4) nascent ring canal formation. Consistent with this, we noted that 3.9 % of Pav-labeled structures in dividing cells were midbodies in fixed preparations; the remaining 94.1% of structures were ring canals (n=439). In contrast, the fluorescence intensity of presumed Pav::mCherry-labeled midbody remnants that result from completed cytokinesis in the somatic hub cells remained static (Fig. 1E, 1F). This indicates that the reduction in midbody signal in dividing germline cells indeed reflects a change in Pav levels at the ring canal and was not due to photobleaching. Furthermore, imaging of photoconverted Pav::Dendra2 (driven by *nanos*-Gal4) in dividing germ cells recapitulated the changes in Pav fluorescence intensity during ring canal formation and revealed that ring canal formation is the direct result of midbody reorganization and occurs in the absence of new protein synthesis (Extended Data Fig. 2). We also examined the behavior of the binding partner of Pav, the RacGAP subunit of the centralspindlin complex, called Tumbleweed (Tum) in *Drosophila*. We found that GFP::Tum exhibited similar changes in fluorescence intensity over the course of ring canal formation (Extended Data Fig. 3) indicating that the entire centralspindlin complex exhibits dramatic changes in localization and fluorescence intensity. These data indicate that ring canal formation in the testis occurs via the formation of a midbody intermediate followed by distinct changes in centralspindlin levels and organization.

Evidence supporting the model of contractile ring arrest in the formation of ring canals comes from studies in the *Drosophila* female germline: Interference of non-muscle Myosin II dephosphorylation through mutations in the myosin-binding subunit of myosin light chain phosphatase, which negatively regulates Myosin II-mediated contraction, resulted in small ring canals that were attributed to over-constriction of the contractile ring (18). This study suggested that dephosphorylation of Myosin II is necessary to arrest the contractile ring resulting in incomplete cytokinesis and was the first genetic evidence implicating proteins necessary for contractile ring constriction in ring canal formation. However, the phenotype was specific to the female germline raising the question of whether different mechanisms drive ring canal formation during spermatogenesis and oogenesis. To address this difference, we examined Pav behavior during *Drosophila* oogenesis. We performed live imaging of Pav::GFP in the germarium, where incomplete mitosis occurs, and found that ring canals form following reorganization of Pav-labeled germline midbodies as in the testis (Fig. 2A, Extended Data Fig. 3). The average time from anaphase onset to ring canal formation was 50 minutes (n=3), which was slower than in the testis and likely the result of differences in cell cycle lengths. Consistent with this, we found fewer dividing ovarian cells with midbodies; only 1.2% of Pav structures in dividing cells were midbodies (n=250). Germline midbodies were present during all four mitotic divisions. Consistent with previous quantifications of ring canal size (22), we found that midbody diameter and nascent ring canal size was smaller with each successive division as cysts moved posteriorly in the germarium (Extended Data Fig. 1).

**Fig. 2.**
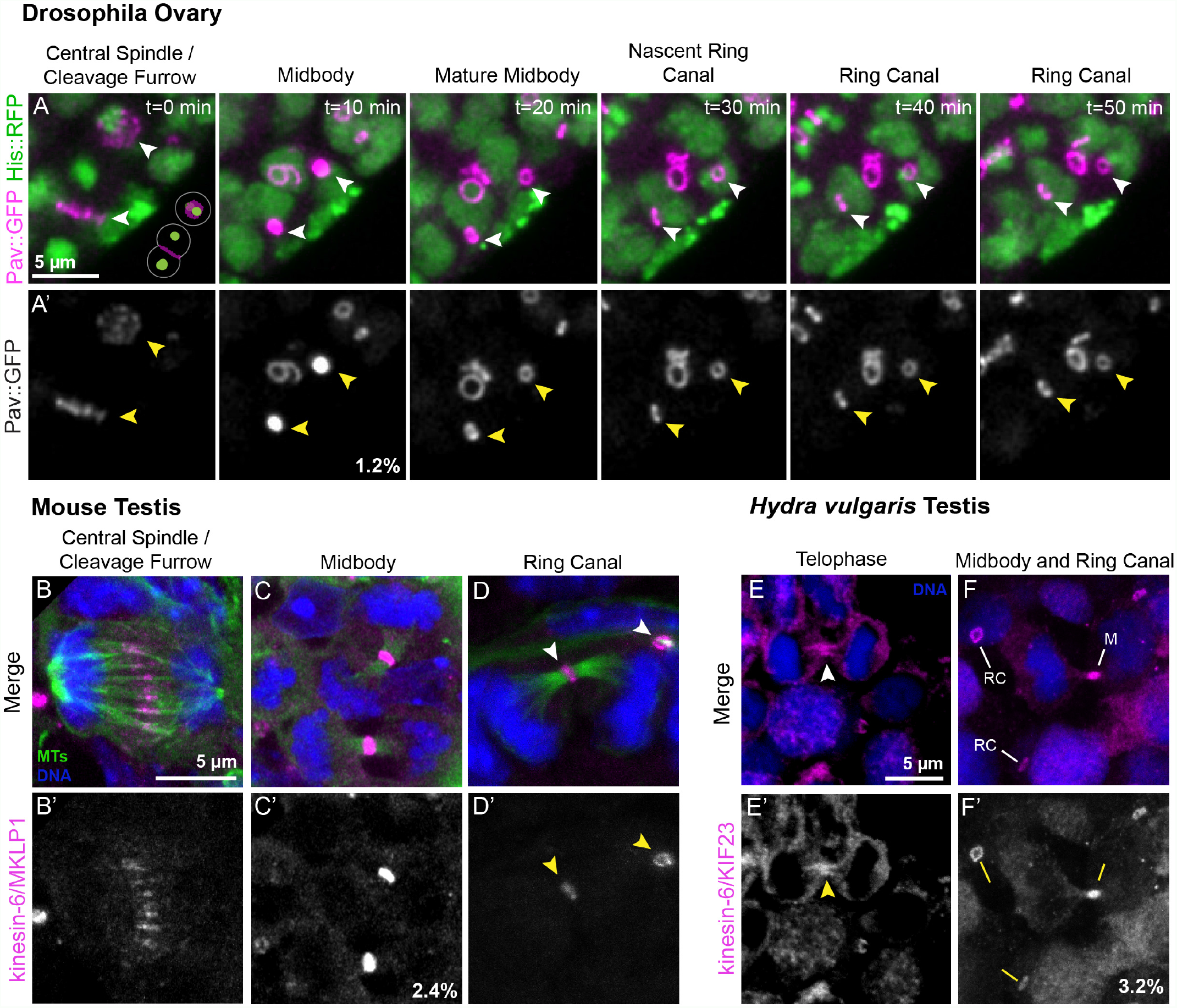
Germline midbody intermediates are a conserved feature of ring canal formation. (A) Stills from a time-lapse movie of a germarium expressing Pav::GFP and Histone::RFP. In the plane of focus are two actively dividing cells of a four-cell cyst; the planes of division are oriented perpendicular to each other as depicted in the schematic at the lower right of t=0 minutes. Arrowheads indicate the progression of cleavage furrows to ring canals. 1.2% of Pav-labeled structures are midbodies. B) Mouse testis sections were fixed and stained to detect microtubules (MTs, green), MKLP1 (magenta), and DNA (blue). (B-B’) A dividing germ cell at anaphase. (C-C’) MKLP1-labeled midbodies are present at the intercellular bridge connected segregated sister chromatids; 2.4% (n=242) of MKLP1-positive structures are midbodies. (D-D’) Nascent ring canals (arrowheads) are marked by the association of open ring canal with luminal microtubules. (E-F’) *Hydra* testes were fixed and stained to detect KIF23 (magenta) and DNA (blue). (E-E’) A dividing spermatocyte with enrichment of KIF23 at the intercellular bridge (arrowhead). (F-F’) Ring canals (“RC”) and a midbody (“M”) are present in the field of view; 3.2% (n=572) of KIF23-labeled structures in dividing cells appeared as midbodies

Together, our data support a mechanism in which midbody formation rather than contractile ring arrest precedes ring canal formation during oogenesis and suggest a common mechanism of ring canal formation during *Drosophila* gametogenesis in males and females.

### Germline midbody intermediates are a conserved feature of ring canal formation

We sought to investigate whether the reorganization of germline midbodies represents an evolutionarily conserved feature of ring canal biogenesis by examining ring canal formation in the male germlines of distantly related animal species, mice and *Hydra*. In mice, there is some evidence that midbodies are involved with ring canal formation. Examination of spermatogenesis in *tex14-/-*mice revealed the presence of MKLP1-labeled midbody-like foci (16). However, it was not clear whether midbody foci are a normal step in the ring canal pathway. Therefore, we carefully examined the localization of MKLP1 (Pav ortholog) and tubulin at the mitotic intercellular bridges in testes from postnatal day 8 mice (Fig. 2B-D’). The majority of MKLP1 signal was found at ring canals (76.4% (n=242); Fig. 2D), as expected. However, we also observed MKLP1-labeled midbodies (2.4% of all MKLP1-positive structures; Fig. 2C), providing evidence that germline midbodies form during mouse cell divisions leading to ring canals. Additionally, we found that the levels of MKLP1 appeared to decline as ring canals formed (compare Fig. 2C’ to 2D’), as we observed in *Drosophila*. These data confirm the presence of germline midbodies in the mouse testis and support the conclusion that a midbody-to-ring canal transition is conserved among bilaterians.

To determine if this behavior of germline midbodies is conserved outside of bilaterians, we examined the localization of the MKLP1/kinesin-6 ortholog KIF23 in testes of the cnidarian *Hydra vulgaris. Hydra* share a common ancestor with the phylum Bilateria, and spermatogenesis takes place in an anatomically simple testis without additional accessory structures like those found in more recently evolved metazoans (23). *Hydra* ring canals have been documented in electron microscopy studies, but the identity of ring canal proteins was unknown. Examination of single cell sequencing datasets revealed that *kif23* transcripts are enriched in male germ cell and nematocyst cell lineages (24), cell types known to have intercellular bridges (4). To study KIF23 localization, we generated and validated a polyclonal antibody against KIF23 amino acids 208-686 (Extended Data Fig. 4), a region that was previously used to generate an antibody to *Drosophila* Pav (20). We stained *Hydra* testes with anti-KIF23 and found that 3.2% of KIF23-labeled structures (n=572) were germline midbodies that localized to the cytoplasmic bridge connecting dividing cells (Fig. 2E-F’) and the majority of remaining KIF23-localized to mature ring canals. Thus, as in the *Drosophila* and mouse germlines, midbodies in the *Hydra* testis were few relative to the number of open ring canals.

From these data, we conclude that the formation of ring canals from germline midbodies is an evolutionarily conserved process and that the midbody-to-ring canal transition represents a key step in ring canal biogenesis.

### Midbody reorganization and stabilization of the midbody ring facilitates ring canal formation

To further characterize the germline midbody-to-ring canal transition, we examined known midbody ring and ring canal components in the *Drosophila* testis using live imaging. Ring canals in the testis have a Septin-rich cytoskeleton made of Septins 1, 2 and Peanut (Septin 7). We imaged Septin 2 (Sep2::GFP) as a representative Septin subunit (Fig. 3A-A”). During ring canal formation, Sep2::GFP localized to the cleavage furrow with Pav::mCherry, but not to the central spindle, forming a ring around the central spindle population of Pav (Fig. 3A-A”, t=0 minutes). Following the formation of the Pav-containing midbody focus, Sep2::GFP was present as a ring around the midbody similar to its localization to the midbody ring in cells that normally undergo complete cytokinesis (25) (Fig. 3A-A”, t=6 minutes). However, rather than progressively disassembling as has been shown in completely dividing cells (26), Sep2::GFP became co-localized with Pav::mCherry in the nascent ring canal (Fig. 3A-A”, t=30 minutes). These data reveal differences in the dynamics of known cytokinesis proteins during incomplete cytokinesis and ring canal formation and suggest that stabilizing factors prevent midbody disassembly or degradation.

**Fig. 3.**
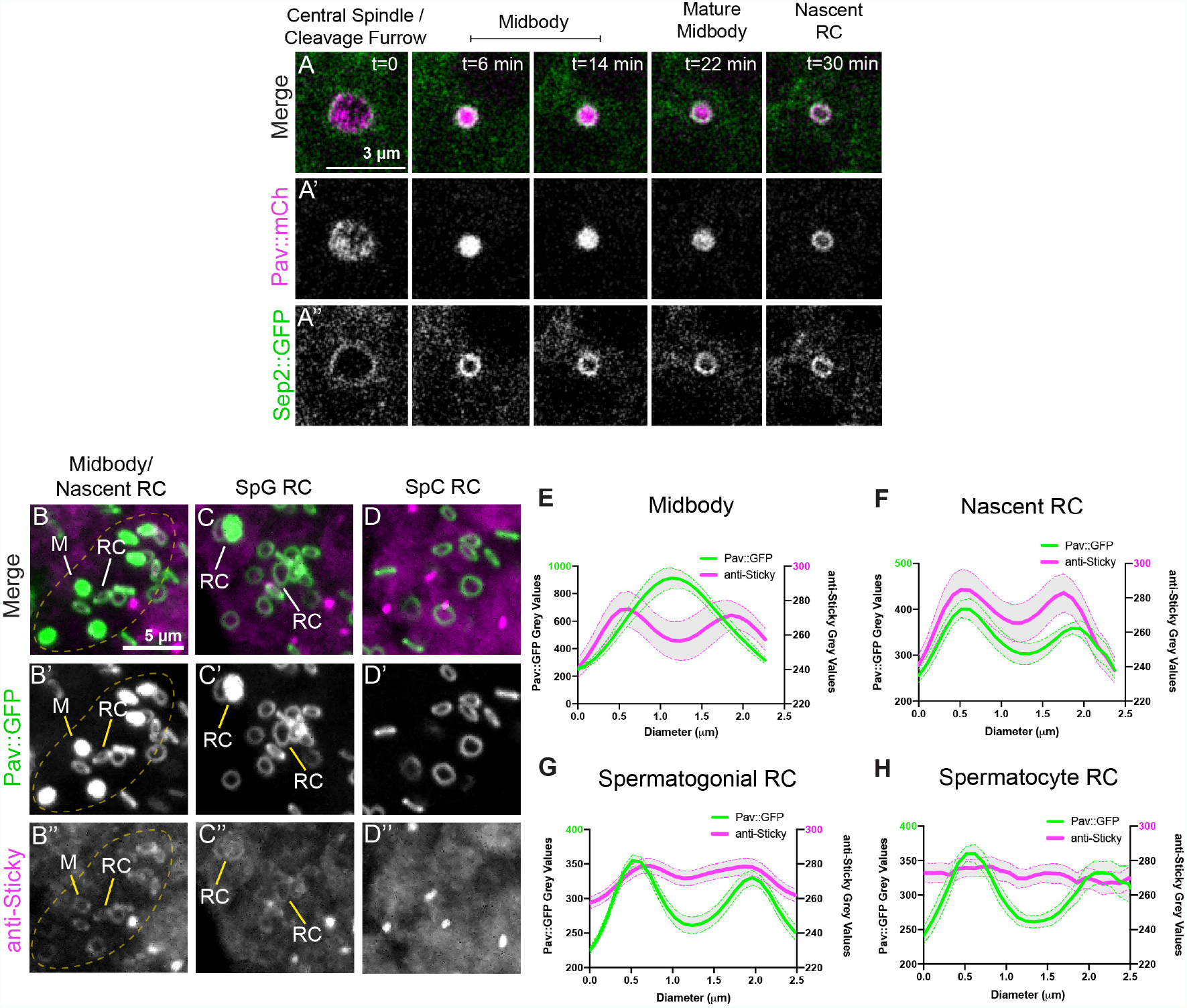
*Drosophila* ring canal formation results from reorganization of germline midbodies. (A-A”) Live imaging of Sep2::GFP and Pav::mCh in the *Drosophila* testis. Sep2::GFP localizes to the cleavage furrow and midbody ring around the Pav-labeled midbody. Sep2::GFP remains in a ring during the midbody-to-ring canal transition and is co-localized with Pav::mCh in the nascent ring canal. (B-D”) Pav::GFP-expressing testes were fixed and stained for Sticky. (B) Outlined in the dashed yellow line is a presumed 16-cell germline cyst, marked by 6 Pav::GFP midbodies (2 are out of the field of view) and 7 nascent ring canals. All Pav midbodies have a ring of Sticky (marked by “M”); all nascent ring canals (“RC”) show co-localization of Pav and Sticky proteins. (C) Ring canals present in spermatogonial (SpG) cells are positive for Sticky. (D) Sticky is absent from ring canals in spermatocytes (SpC). (E-H) Pav::GFP fluorescence intensity (green trace, left Y axis) versus Sticky (magenta trace, right Y axis) is plotted during the midbody (n=9), nascent ring canal (n=5), spermatogonial ring canals (n=20) and spermatocyte ring canals (n=13); Thick lines are the means and error bars (shaded grey) reflect the SEM.

We next examined the localization of the midbody ring protein Sticky/Citron kinase (Citk). Citk is required for the stereotypic organization of proteins within the midbody during late cytokinesis and is indirectly required for cytokinetic abscission (27–29). Interestingly, mutations in Citk cause male sterility in mice due to the loss of intercellular germline bridges as a consequence of complete cytokinesis (30). In *Drosophila* spermatogenesis, mutations in *sticky* result in an increase in multi-nucleate spermatocytes due to failed cytokinesis as well as smaller than average ring canals, attributed to over-constriction of the cleavage furrow (31). Immunolabeling of Sticky in fixed *Drosophila* testes expressing Pav::GFP revealed that Sticky localized to the midbody ring and nascent ring canal (Fig. 3B-B”, 3E, 3F). Sticky was present at reduced levels on nascent ring canals in spermatogonia (SpG) where it co-localized with Pav::GFP (Fig. 3C-C”, 3G), and was absent from mature ring canals in primary spermatocytes (SpC) (Fig. 3D-D”, 3H). The transient localization of Sticky to the mid-body ring and nascent ring canals suggests a possible function of Sticky to facilitate the midbody-to-ring canal transition.

### Citron kinase/*Drosophila* Sticky is required for the midbody– to-ring canal transition

To investigate the role of Sticky during ring canal formation, we examined Pav::GFP in testes with *sticky* expression reduced by RNAi. We labeled tissues with Hts/Adducin antibodies to visualize the fusome, a branched organelle that extends through all ring canals in germline cysts, to provide another metric of proper ring canal formation (Fig. 4A-B’, 4C-D’). In control testes, ring canals were uniform in size with an average diameter of 2 *µ*m and a continuous fusome filled the ring canal lumens (Fig. 4A-B”, 4F). In contrast, ring canals in *nos>sti RNAi* testes were significantly smaller, as previously described (31), with an average ring canal diameter of 1.1 *µ*m (Fig. 4C-D”, 4F). Interestingly, we observed that the ring canal phenotype was not restricted to spermatocytes, as previously described by Naim et al. (2004), but manifested in mitotically dividing cells demonstrating a general requirement for Sticky in ring canal formation at all stages of spermatogenesis. In addition, 31.4% of Pav::GFP structures were focal, appearing as persistent midbodies (Fig. 4C-D”, 4E, 4G), similar in morphology of those found in the somatic hub. At these focal sites, the fusome was either discontinuous with obvious gaps or a thread-like structure, indicative of perturbed or complete cytokinesis (Fig. 4C-D”). To confirm that the Pav midbody-like foci represent persistent midbodies, we performed live imaging of Pav::GFP in *nos>sti RNAi* testes (Fig. 4E). In *nos* control testes, the time to ring canal formation was an average of 26.7 minutes (± 2.3 minutes, SEM) (Fig. 4H). Of the ring canals that formed successfully in *nos>sti RNAi* testes, the average time to ring canal formation was 37.4 minutes (± 1.6 minutes, SEM). However, many midbodies failed to resolve into ring canals over the course of our time-lapse imaging (Fig. 4E). Additionally, we observed that *nos>sti RNAi* males had reduced fecundity (Fig. 4I). These data demonstrate a requirement for Sticky in the midbody-to-ring canal transition and, taken together with the fecundity phenotype, suggest an important role for midbody reorganization in the formation of ring canals and fertility.

**Fig. 4.**
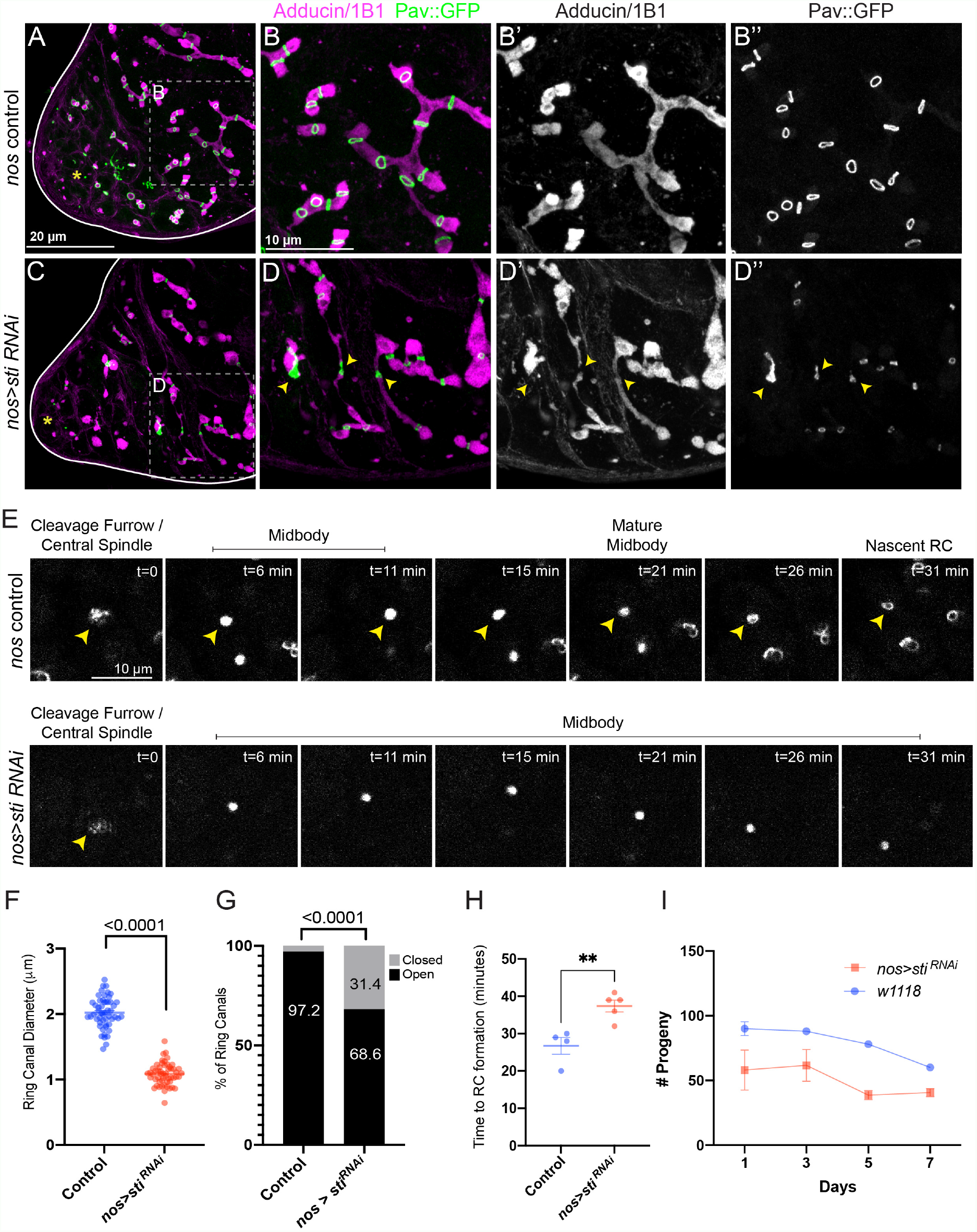
Sticky facilitates the midbody-to-ring canal transition. (A-B”) Control Pav::GFP testes were fixed and stained for Adducin/1B1 (magenta). The apical tip is marked by a yellow asterisk. In controls, the fusome (labeled with Adducin/1B1) is continuous and connects all ring canals in a germline cyst. Ring canals are open. (C-D”) *nos>sti RNAi* testes. Yellow arrowheads mark closed ring canals and points where fusome structure is perturbed. (E) Live imaging of Pav::GFP in control and *nos>sti RNAi* testes. In *nos>sti RNAi* testes, midbody foci form but fail to resolve into ring canals in the same time frame as controls. (F) In *nos>sti RNAi* testes where the ring canal diameter can be scored (control n=52; *nos>sti RNAi* n=54), ring canal diameter is reduced (1 *µ*m ± 0.02, SEM) as compared to controls (2 *µ*m ± 0.03, SEM). (G) *nos>sti RNAi* testes have a higher percentage of foci (“closed”) (n=116/369) as compared to controls (n=2/73) (p=<0.0001, fisher’s exact test). (H) Of the ring canals that form in *nos>sti RNAi* testes, the time to ring canal formation is delayed (p=0.0054). (I) *nos>sti RNAi* males are also sub-fertile compared to controls (p=0.0063, paired two-tailed t-test).

### Abscission inhibition occurs at the germline midbody

The absence of abscission during incomplete cytokinesis suggests that the germline midbody functions differently from the mid-bodies formed during complete cytokinesis. To better understand the germline midbody, we examined the behavior of microtubules during ring canal formation as a readout of mid-body behavior. We performed live imaging of Pav::mCherry and a GFP-tagged microtubule-binding protein Jupiter::GFP (32, 33) to evaluate microtubule behavior. We found that microtubules persisted throughout incomplete cytokinesis and ring canal formation, rather than undergoing severing or de-polymerization as documented during abscission (Fig. 5A-A”). Nascent ring canals contained microtubules for an extended time within the ring canal lumen (Fig. 5A-A”, t=67 minutes) before they ultimately disappeared from the lumens of older ring canals. We also observed persistent microtubules in the lumens of nascent ring canals in the mouse testis (Fig. 2D). If depolymerization of microtubules is a pre-requisite for ab-scission, then microtubule persistence may contribute to the absence of abscission in the germline.

**Fig. 5.**
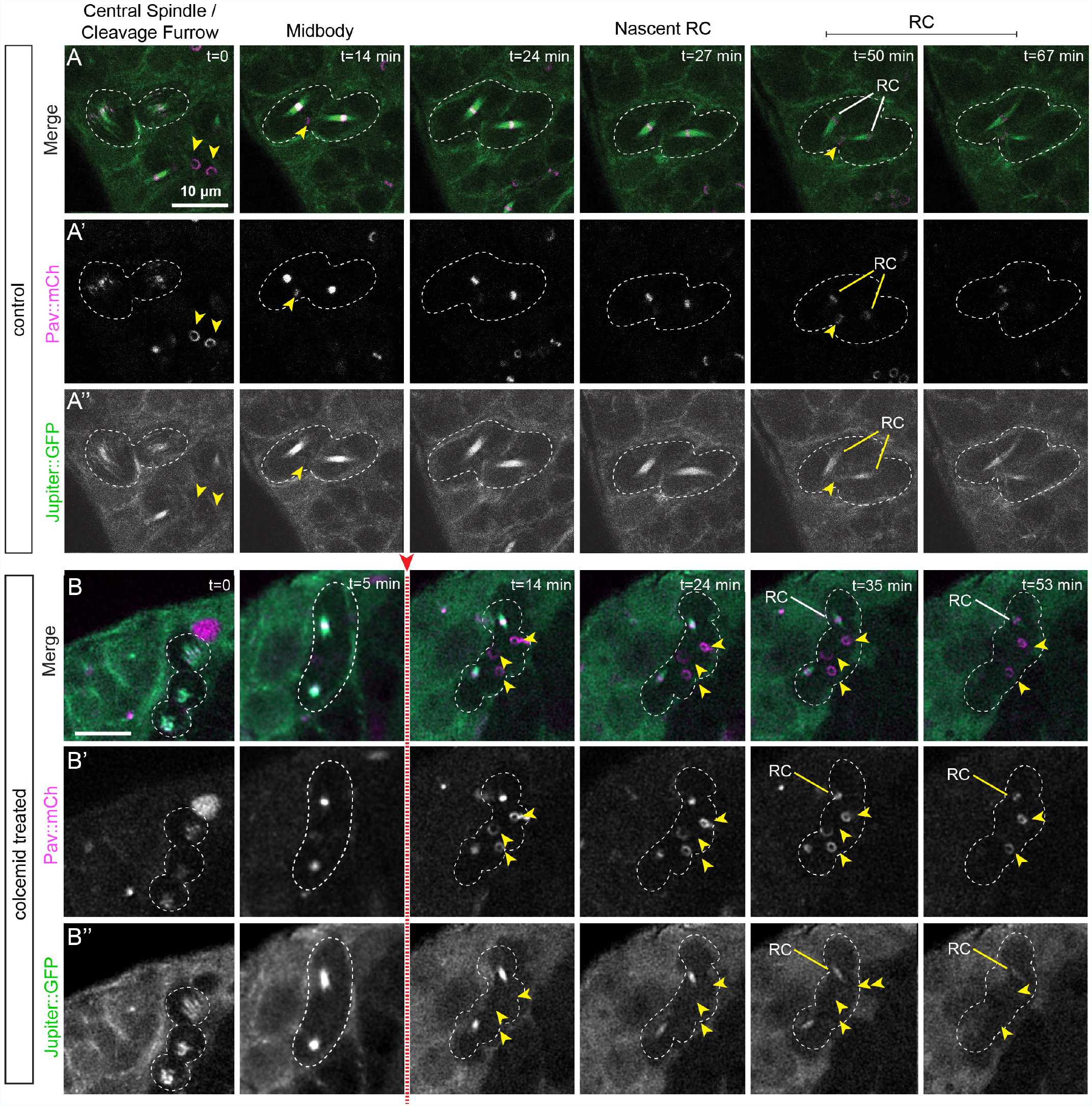
Stable intercellular bridge microtubules are maintained during ring canal formation. (A) Pav::mCh was imaged with Jupiter::GFP to visualize microtubules. An actively dividing two-cell cyst (outlined) with single, mature ring canal (arrowhead in t=14 minutes frame) will generate two new ring canals (labeled “RC” at t=50 minutes) that maintain Jupiter::GFP-labeled microtubules in the lumen. In contrast, two mature ring canals (arrowheads in t=0 minutes frame) do not contain luminal Jupiter::GFP signal revealing the microtubules, while present during the midbody-to-ring canal transition, are eventually removed in the mature ring canal. (B) Testes expressing Pav::mCh and Jupiter::GFP were treated with 100 *µ*M colcemid following formation of the nascent midbody (red arrow). A dividing four-cell cyst (outlined) with mature ring canals (arrowhead in t=14 minutes frame) will divide to generate an eight-cell cyst and generate four new ring canals (only one of which is in frame at t=53 minutes). Following colcemid treatment, cytoplasmic microtubules are disrupted but the spindle midzone microtubules remain stably associated with the midbody and nascent ring canal and there are no obvious defects in ring canal morphology.

In completely dividing cells, midzone microtubules are stable and insensitive to disruption by microtubule depolymerizing drugs (34, 35). To ascertain whether the spindle midzones during incomplete cytokinesis are inherently different from the midzones described during complete cytokinesis, we treated testes expressing Pav::mCherry and Jupiter::GFP with the mi-crotubule depolymerizing drug colcemid. Colcemid treatment resulted in a rapid and complete destabilization of cytoplasmic and centrosomal microtubules but did not appreciably affect the midbody microtubules (Fig. 5B-B’). These data revealed that the midzone population is no less stable than midzones described during complete cytokinesis, but do not discount the possibility that spindle midzones in the germline are more stable than their somatic counterparts. In fact, hyper-stability of microtubule midzones in mammalian cells has been shown to result in abscission failure, with cells connected by long intercellular bridges (36). However, we were neither able to enhance microtubule stability nor fully disrupt the midzone microtubules to directly test the effect on ring canal formation.

During cytokinesis in completely dividing cells, central-spindlin at the midbody links to the abscission machinery through a series of direct protein-protein interactions: In mammals, centralspindlin binds to the dimeric coiled-coil protein CEP-55 that in turn interacts with both Alix (Alg-2 interacting protein) and the ESCRT-I subunit Tsg101 that then recruit the ESCRT-III complex required for abscission (37–39). ESCRT-III complex subunits recruit the microtubule severing complex Spastin, which promotes disassembly of microtubules in the midbody in preparation for abscission (40). In *Drosophila*, which lack a CEP-55 ortholog, the ESCRT-associated protein Alix binds Pav directly (41). We investigated the localization of abscission pathway proteins involved with microtubule severing and membrane abscission (Extended Data Fig. 5A). Consistent with our observation of persistent microtubules, we did not detect Spastin::GFP at midbodies or nascent ring canals even when over-expressed (Extended Data Fig. 5B). Similarly, we did not detect ESCRT-III/Shrub::GFP *(nos>UAS-shrub::GFP)* at germline midbodies or nascent ring canals (data not shown). However, we did find that Alix::GFP localized to germline midbodies, similar to its localization to the midbodies during complete cytokinesis in *Drosophila* female germline stem cells (42), and in nascent ring canals (Extended Data Fig. 5C, 5D), possibly by direct interaction with Pav as previously shown (41).

As another readout of abscission progression, we examined the behavior of F-actin. During midbody maturation in completely dividing cells, F-actin is gradually removed from the cleavage furrow in preparation for abscission, and perturbation of F-actin remodeling factors can delay cytokinesis (43, 44). Fixed preparations of testes labeled with phalloidin as well as live imaging of F-actin with LifeAct::mRuby revealed that contractile ring F-actin was mostly disassembled at the germline midbody and absent from mature ring canals, consistent with previous studies that have demonstrated that mature *Drosophila* male ring canals do not contain F-actin filaments (17, 45) (Extended Data Fig. 5E-G”).

Taken together, these data demonstrate that canonical midbody function is altered at germline midbodies such that key abscission pathway components do not accumulate. Our data also raise the question of whether stable microtubules play a role in the transition from midbody to open ring canal.

## Discussion

Until now, studies of ring canal formation have lacked the spatiotemporal resolution required to understand the pathway of ring canal biogenesis. Our results reveal the temporal dynamics of incomplete cytokinesis and ring canal formation for the first time. We provide strong evidence that germline midbodies function differently in cells that make ring canals, and discover that germline ring canal formation occurs via an evolutionarily conserved germline midbody intermediate.

Our live imaging data in the *Drosophila* germlines challenge previously held ideas of contractile ring arrest and inform a new model that is consistent with observations of germline midbodies during spermatogenesis in mice. Following midbody formation in incomplete cytokinesis, the midbody reorganizes to form a midbody ring that is stabilized and is transformed into a ring canal (Fig. 6). Our data implicate the evolutionarily conserved centralspindlin complex as a driver of ring canal formation, as it undergoes a stereotypic relocalization that precedes ring canal lumen formation. We identify a role for the Citron kinase ortholog Sticky during the midbody-to-ring canal transition in *Drosophila* males that may contribute to midbody reorganization in other species and cell contexts.

**Fig. 6.**
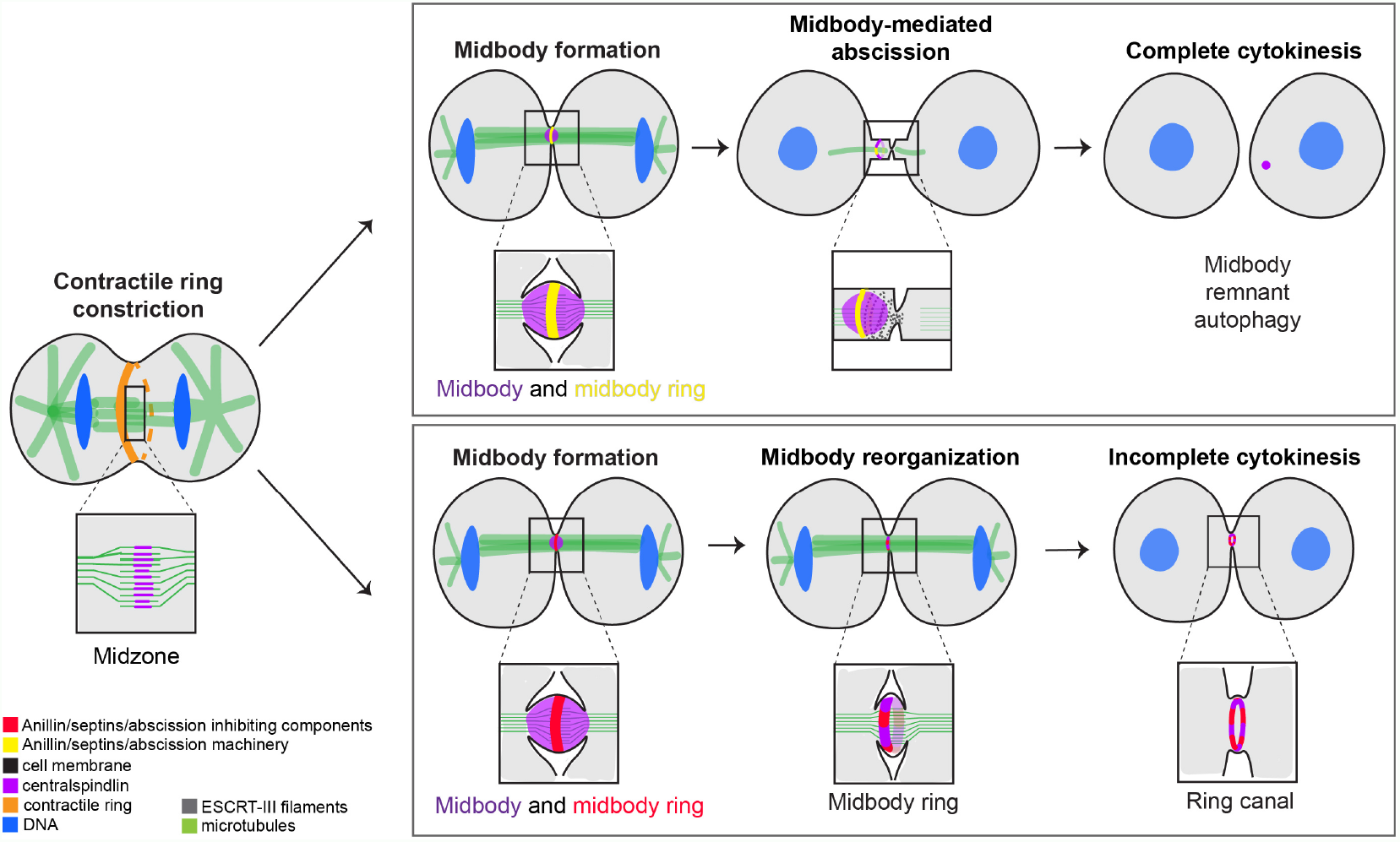
Incomplete and complete cytokinesis pathways diverge following midbody formation. Following anaphase onset, a central spindle/spindle midzone assembles between the segregating chromosomes and an actomyosin contractile ring assembles at the cell cortex. As the contractile ring constricts, proteins in the central spindle are compacted resulting in the formation of a midbody (purple) and midbody ring (yellow). Proteins necessary to carry out abscission are recruited to the midbody and incorporated in the midbody ring. Abscission machinery components promote microtubule depolymerization and abscission via the formation of ESCRT-III filaments (dotted grey spirals). During incomplete cytokinesis, we propose that molecules necessary for abscission inhibition and midbody reorganization are recruited to the midbody and/or midbody ring (depicted as a red ring) promoting ring canal formation. Microtubules are disassembled much later in a manner independent of Spastin severing activity.

### Germline midbody formation is an intrinsic feature of game-togenesis

Our data support the view that germ cells use a shared mechanism of ring canal formation and that evolutionarily shared components drive the formation of ring canals. Contrary to previous studies of ring canal formation that have relied on fixed immunofluorescence assays (17, 18, 46), our live imaging of conserved ring canal components in the *Drosophila* male and female germlines has revealed that germline midbody formation, not contractile ring arrest, is a shared feature of gametogenesis. Our investigation of germline midbody formation in distantly related species reveals a mechanism of incomplete cytokinesis giving rise to ring canals that is conserved between the sexes and across evolution. Surprisingly, through our characterization of ring canal formation, we discover that incomplete cytokinesis shares more features with complete cytokinesis than previously appreciated. Rather than the pathways of cytokinesis diverging shortly after cleavage furrow formation and ingression, our data suggests that both cytokinesis programs assemble a midbody that appears to act as a pause point, allowing the cells to either recruit components to separate newly formed daughter cells, or initiate a mechanism that prevents abscission and promotes formation of cells in a syncytium.

### Midbodies function differently in the germline

Although mid-bodies form in both completely and incompletely dividing cells, we show that midbody function differs between the cytokinetic programs, suggesting that midbody formation is the point at which the pathways of cytokinesis diverge (Fig. 6). During complete cytokinesis, abscission pathway components localize to the midbody in preparation for membrane severing (43). We did not detect abscission pathway components at the midbody in germline cells, nor did we observe evidence of canonical microtubule severing, suggesting that the absence of abscission in the germline is likely due to the absence of abscission proteins at the midbody. One hypothesis is that abscission inhibition factors, functionally analogous to TEX14 in mammals, reside at the germline midbody and prevent the recruitment of downstream ESCRT-associated proteins. Interestingly, when TEX14 is over-expressed in HEK293T cells, TEX14 localizes to the midbody and prevents abscission (15) supporting our hypothesis that abscission inhibition factors in *Drosophila* reside at the midbody.

Alternatively, differences in midbody function could relate to the lifetime of the germline midbody during ring canal formation. In completely dividing cells, cleavage furrow ingression occurs in less than one hour and is followed by the longer midbody-mediated process of abscission that lasts several hours (29). We observed that germline midbodies persist for 20-30 minutes suggesting that their short lifetime may contribute to abscission inhibition. Rapid reorganization of the germline midbody into a ring may prevent recruitment of proteins that would otherwise localize to the midbody during the prolonged midbody maturation process that precedes abscission of nascent daughter cells. Whereas the timing of ring canal formation during mouse spermatogenesis has not been determined, it is reasonable to consider that the timing of ring canal formation is longer as the timing of spermatogenesis is much longer than in *Drosophila* (34.5 days (47) versus 10.4 days (48)), respectively. It will be interesting to correlate the timing of ring canal formation in different species with the presence or absence of TEX14 to determine if the transient nature of the germline midbody is sufficient to prevent abscission.

### Ring canals represent persistent midbody rings

Our data reveals that centralspindlin in the midbody reorganizes to join proteins in the midbody ring suggesting that ring canals form from stabilized midbody rings (Fig. 6). The mechanism underlying reorganization of centralspindlin from its midbody localization to that in the midbody ring is unknown, but our live imaging data suggests some hypotheses that warrant further investigation. As centralspindlin relocalizes from the midbody to the ring canal, protein in the center of the focus is removed, either via degradation or relocalization, leaving behind a centralspindlin ring that colocalizes with the midbody ring protein Citron kinase/Sticky and the septins. It is possible that during the transition, the microtubule motor domain of MKLP1 releases the microtubules that comprise the spindle midzone and, in doing so, is reorganized to form a ring that encircles the microtubule bundle. Alternatively, there may be two sub-populations of centralspindlin within the midbody, i) the cortically associated ring population that is protected from degradation, and ii) the spindle midzone associated population that is selectively degraded during the midbody-to-midbody ring/ring canal transition. Interestingly, microtubule spindles in *Drosophila* primary spermatocytes are comprised of two microtubule populations, an “interior” central spindle and a “peripheral” astral microtubule population, the former of which is associated with the microtubule associated protein Orbit/Mast/CLASP (49). Therefore, one possibility in our model is that centralspindlin bound to the “interior” microtubule population, perhaps via interactions with Orbit/Mast/CLASP, is targeted for degradation leaving behind only the cortically associated ring. Following midbody reorganization, the ring canal is stabilized and protected from degradation. It is known that ring canals are rich in phosphotyrosine epitopes and glycoproteins, but whether these post-translational modifications protect the ring canal from autophagic degradation is unclear and an area of future research.

### Ring canal formation is more than just abscission inhibition

It is unlikely that abscission inhibition alone is sufficient to produce a ring canal. In dividing *Drosophila* germline stem cells, the abscission factor Alix localizes to the midbody and promotes abscission via direct interaction with ESCRT-III/Shrub (42). However, *alix* mutant germline stem cell divisions often fail to undergo abscission and daughter cells remain inter-connected by persistent midbody rings demonstrating that midbody ring formation proceeds in the absence of abscission (42). The effect of blocking abscission in somatic cells is quite different. Knockdown of Shrub in *Drosophila* S2 cells results in chains of cells connected by midbody-like foci (50) demonstrating that abscission inhibition alone does not result in midbody reorganization and ring canal formation. These contradictory results suggest the possibility that cell type-specific factors are required to reorganize the midbody to an open ring canal when abscission is inhibited. Incomplete cytokinesis is therefore the likely result of two coordinated pathways: abscission inhibition and midbody reorganization.

### Citron kinase may coordinate the pathways of abscission in-hibition and midbody reorganization

In addition to central-spindlin, the evolutionarily conserved serine/threonine kinase Citron kinase/Sticky is an important player in ring canal biogenesis. Whereas studies of Citron kinase function in completely dividing cells reveal critical roles in promoting abscission, our analyses in the *Drosophila* germline reveal an important role for Sticky in preventing abscission. In testes depleted of Sticky protein, we observe a significant percentage of smaller than average germline midbodies that are coincident with complete cytokinesis events in the germline. Additionally, we have identified a new role for Citron kinase/Sticky in promoting the timely reorganization of germline midbodies to ring canals. Together, these data suggest that, not only is Sticky required for the proper architecture and size of the germline midbody -perhaps functioning in an analogous manner as in completely dividing cells-but that Sticky is also required to reorganize centralspindlin during the midbody-to-ring canal transition. Our observations in the *Drosophila* germline, combined with observations of aberrant abscission in the mouse male germline (30), are hard to reconcile with the described phenotype of Citron kinase in completely dividing cells where depletion or mutation in Citron kinase leads to abscission failure (28, 51–55). How can Citron kinase have different cytokinesis functions? One possible explanation is that Citron kinase, as a scaffolding protein, may interact with different suites of proteins at the germline versus somatic midbody to effect cell division. Alternatively, Citron kinase phosphorylation of cell-type specific targets may differentially affect protein function, activity, or binding partners. Considering the conservation of Citron kinase, it is tempting to speculate that Citron kinase, together with centralspindlin, may act as part of a conserved pathway of ring canal formation to coordinate midbody reorganization with abscission inhibition.

### Do ring canals reflect the evolution of multicellularity?

Our findings in distantly related species underline the evolutionary conservation of germline midbody formation during incomplete cytokinesis that results in ring canals and reveals interesting insights about the evolution of cell division processes. During complete cytokinesis, centralspindlin marks the midbody core that remains associated with the membrane and is inherited by one of the nascent daughter cells as a midbody remnant. However, following abscission the midbody remnant undergoes autophagy. In contrast, the ring canal remains stably associated with the membrane and serves as an important channel for communication between syncytial cells. During certain life cycle stages in single-celled colony-forming eukaryotes like the choanoflagellates, the closest living relatives of animals, and in the green algae Volvox, neighboring cells in colonies are connected by intercellular bridges that appear similar to ring canals (56, 57). It is unclear whether centralspindlin localizes to the intercellular bridges in the colonies; however, given the presence of centralspindlin homologs in these genomes, it is exciting to consider that centralspindlin dynamics during incomplete cytokinesis may have contributed to the ancestral program that produced chains of cells during the evolution of multicellularity. Later embellishments in the form of abscission proteins were introduced to produce completely separated cells and the original function of the midbody ring to form a ring canal was lost.

As incomplete cytokinesis events are observed during normal development and are prevalent in disease states, investigation into the mechanism of ring canal formation is poised to reveal insights into the fundamental mechanism of incomplete cytokinesis across biology.

## Materials and Methods

### Fly husbandry and strains

The following FlyTrap line was used: Jupiter::GFP (CB05190). The following fly strains were obtained from the Bloomington Drosophila Stock Center: *nos-Gal4* (7303), *his2av::mRFP* (23651), *his2av::eGFP* (24163), *sti shRNA* (35392), *sep2::GFP* (26257), *lifeAct::mRuby* (35545), and *sfGFP::tum* (76264). We imaged Pavarotti (Pav) at endogenous levels with Pav::GFP (21) or with Pav::mCherry (this study). *uasp-GFP-alix* flies were obtained from Kaisa Haglund (Oslo University Hospital) and *uas-spas-eGFP* flies from Damien Brünner (University of Zurich).

### Construction of Pav::mCherry

Pav::mCherry was generated in a *w*^1118^ background by replacing the endogenous stop codon of *pav* with a C-terminal mCherry-LoxP-3XP3-eGFP-LoxP knock-in cassette via CRISPR/Cas9-mediated genome editing by homology-dependent repair (performed by WellGenetics, Inc.). Following screening, 3XP3-GFP was removed by crossing to flies expressing Cre recombinase.

### Construction of Pav::Dendra2

The coding sequence for Dendra2 was amplified from genomic DNA of *–*-tubulin 84B::Dendra2 flies (51315). Full-length Pav cDNA and Dendra2 were cloned into the pUASz vector with Gibson Assembly (NEB). Pav fused to Dendra2 is spaced with the flexible linker GGSGGSGGSG. The final plasmid was inserted at the attP40 site on 2L by Genetivision (Houston, TX).

### Live imaging in *Drosophila* testes

Live imaging of ring canal formation in Drosophila testes was performed as previously described (58). Newly eclosed males were dissected in Ringer’s solution (200 mM NaCl, 4 mM KCl, 0.3 mM CaCl2 2H20, 5 mM NaHC03, 0.13 mM NaH2PO4 H20) and then transferred in Ringer’s solution to a Poly-L-Lysine-coated coverslip bottom of a round imaging dish (MatTek). Ringer’s solution was removed, allowing the testes to settle on the coverslip, and replaced with imaging medium (15% Fetal Bovine Serum, 0.5X penicillin/streptomycin, 0.2 mg/ml insulin in Schneider’s insect medium). Testes were imaged on either a Bruker Opterra II Swept Field or Zeiss 980 laser scanning confocal microscope. Images were acquired with a Z step size of 1 *µ*m using either a 60X water immersion objective lens (Bruker Opterra II, 1.3 NA) or W Plan-Apochromat 40X dipping lens (Zeiss, 1.0 NA). Images were collected at either 1 minute or 2 minutes intervals. The average Z-distance traveled was 47 *µ*m per time point. Due to contractions of the testis muscle sheath, figures were created from, and analyses were performed on individual Z-sections. When possible, samples were imaged in a temperature-controlled chamber set to 25°C to prevent thermal drift.

### Photoconversion of *nos>pav::dendra2*

Testes from *nos>pav::dendra2* flies were dissected in Ringer’s solution and prepared for live imaging as described above. Photoconversion experiments were carried out on a Bruker Opterra II Swept Field confocal microscope with PlanApo 40X water immersion objective with 1.6X optical zoom (1.15 NA). A single Z-stack was captured prior to photoconversion. A 10 *µ*m^2^ region of interest in a single Z-plane was photoconverted with 50 iterations of 405 nm wavelength, a 50 *µ*s pixel dwell time, and 390 nm pixel size (512 × 512 pixels). Following photoconversion, a Z-stack was acquired every 60 seconds. Green-to-red photoconversion was detected using the same settings.

### Colcemid treatment of *Drosophila* testes

Testes from flies expressing Pav::mCh and Jupiter::GFP were dissected and prepared for imaging as described above. Dividing cells were followed until the formation of a midbody, upon which time 100 *µ*M Colcemid was added to the dish in the medium directly above the testis being imaged. Disruption of cytoplasmic microtubules occured 10 minutes following addition and was evidenced by the loss of Jupiter::GFP-labeled cytoplasmic microtubules.

### Live imaging in *Drosophila* ovaries

Live imaging of ring canal formation in *Drosophila* ovaries was performed as previously described (59). Briefly, newly eclosed females were dissected in Ringer’s solution on a Poly-L-Lysine-coated coverslip of a Mat-Tek imaging dish. Ringer’s solution was removed carefully as to not disrupt the germaria and replaced with imaging medium (15% Fetal Bovine Serum, 0.5X penicillin/streptomycin, 0.2 *µ*g/ml insulin in Schneider’s insect medium). Images were acquired with a Z step size of 0.6 *µ*m using a W Plan-Apochromat 40X dipping lens (Zeiss, 1.0 NA) every 10 minutes.

### *Drosophila* immunocytochemistry in testis

Testes were dissected in Ringer’s solution and fixed for 10 minutes in 4% paraformaldehyde in PBT (phosphate-buffered saline with 0.3% Triton X-100 and 0.5% BSA). Fixed tissue was washed in PBT and permeabilized in 1% Triton X-100 in PBS for 30 minutes at room temperature. Testes were then blocked in for 30 minutes at room temperature and incubated in primary antibody diluted in blocking solution. The following primary antibodies were used: mouse anti-Hts1B1/Adducin (1:50, Developmental Studies Hybridoma Bank) and rabbit anti-Sticky (60) (1:600). Alexa Fluor-conjugated secondary antibodies were used at 1:200 and diluted in blocking solution. Samples were washed in PBT and mounted on slides in ProLong Gold antifade reagent (Invitrogen). DNA was visualized with DAPI (1 *µ*g/ml). Samples were imaged with a Plan-Apochromat 63X 1.4 NA oil immersion objective on Zeiss 980 laser scanning confocal microscope with Airyscan2. Airyscan2 images were processed post-acquisition with the Zeiss Airyscan processing feature.

### Male fertility tests

Fecundity of *sti*-depleted males was assessed by pairing a single male with three *w*^1118^ virgin females. Males were shifted every two days to new vials containing new virgin females for 14 days. The total number of adult progeny was counted to determine fecundity.

### Mouse immunocytochemistry

All mouse work was approved by Yale University’s Institutional Animal Care and Use Committee. Mice were maintained in a barrier facility with 12 hour light/dark cycles. Mouse testes from postnatal day 8 mice were fixed in 4% paraformaldehyde in PBS overnight at 4°C, sucrose-protected in 30% sucrose in PBS overnight at °C, embedded in Optimal Cutting Temperature Compound (Tissue-Tek) prior to freezing, and samples were stored at -80°C. 10-micrometer sections were cut via cryostat. Slides were post-fixed in 4% paraformaldehyde for 8 minutes at room temperature. Sections were washed briefly in PBS and then incubated in a 1% SDS solution for 5 minutes at room temperature in a humid chamber. Sections were washed thoroughly in PBS and then incubated in blocking buffer (3% BSA, 5% Normal Goat Serum, 5% Normal Donkey Serum in PBS-T (PBS containing 0.2% Triton X-100)) for 30 minutes at room temperature. Primary antibodies were diluted in blocking buffer and incubated in a humid chamber for 30 minutes. The following primary antibodies were used: 1:200 rabbit anti-MKLP1 (clone 7C9, Bioss Antibodies, cat. BSM-52401R) and 1:10 mouse anti-alpha Tubulin (clone 4A1, Developmental Studies Hybridoma Bank). After washing in PBS-T, sections were incubated in secondary antibody diluted in blocking buffer for 30 minutes. Alexa Fluor-conjugated antibodies were diluted 1:200 in blocking buffer and incubations were done for 30 minutes at room temperature in a humid chamber. DNA was visualized with DAPI (1 *µ*g/ml). Sections were washed and then mounted in ProLong Gold antifade reagent (Invitrogen). Coverslips were sealed with clear nail polish. Tissue sections were imaged on a Zeiss 980 laser scanning confocal microscope with 63X 1.3 NA oil immersion objective.

### *Hydra* immunocytochemistry

Hydra were relaxed in 2% urethane in Hydra medium for 2 minutes, fixed for 10 minutes in 4% paraformaldehyde in Hydra medium at room temperature, washed 3 times with PBS, permeabilized with 0.5% Triton X-100 in PBS for 15 minutes. Samples were then incubated in blocking buffer (1% BSA, 10% Normal Goat Serum, 0.1% Triton X-100 in PBS) for one hour. Primary antibodies were diluted in blocking buffer and incubated overnight at 4°C. Rabbit Anti-kinesin-6/KIF23 antibody (this study) was used at 1:200. Samples were washed 3 times in PBS and incubated in secondary antibody diluted in blocking buffer for 2 hours at room temperature. Alexa Fluor-conjugated secondary antibodies were used at 1:200. DNA was visualized with 1 *µ*g/ml DAPI. Animals were mounted in ProLong Gold antifade reagent (Invitrogen). Whole mount *Hydra* preparations were imaged on a Zeiss 980 laser scanning confocal microscope with 63X 1.3 NA oil immersion objective.

### Hydra Culturing

*Hydra vulgaris* AEP were asexually cultured at 18°C in Hydra medium by standard procedures (Lenhoff and Brown, 1970) and fed Artemia nauplii 2-3 times a week. Polyps with testes were selected for immunostaining

### *Hydra* KIF23 antibody generation and validation

A fragment of *Hydra* KIF23 (Uniprot T2MND8, GenBank accession number XM 012700539.1) from amino acids 208-686 was cloned into the pET30a (+) expression vector. Recombinant protein was expressed in BL21 StarTM DE3 cells. Purified proteins were used to raise antisera in rabbits (Genscript) from which antibody was then affinity purified. Antibody was verified via Western blot against purified protein that was used as immunogen as well as *Hydra* whole cell lysate. 50 ng purified protein, 3 Hydra equivalents, and 10 *Drosophila* testis equivalents were run on a 4-15% gradient SDS-PAGE gel (BioRad) and transferred at 300 mAmps for 42 minutes to a nitrocellulose membrane (GE Healthcare). The blot was probed with 4 1:5000 anti-KIF23 antibody.

### Image and statistical analysis

Raw image files were adjusted for brightness and contrast in Fiji/ImageJ. Single Z-slices were used for image analyses. Generation of plots and statistical analysis was performed in Graph Pad Prism.

## Acknowledgements

We would like to thank Pier Paolo D’Avino for the anti-Sticky antibody and Kaisa Haglund and Damien Brünner for critical reagents. We thank Celina Juliano for providing *Hydra vulgaris* animals and technical advice. We are also grateful to Kaelyn Sumigray for providing mice and technical advice and Chia-Ling Hsieh for performing mouse testis dissections. Lastly, we would like to thank members of the Cooley and Sumigray labs for helpful discussions as well as Andrew Cox for critical reading of the manuscript. Stocks obtained from the Bloomington Drosophila Stock Center (NIH P40OD018537) were used in this study. We thank the DRSC/TRiP Functional Genomics Resources at Harvard Medical School for providing transgenic RNAi fly stocks used in this study. The Hts/Adducin 1B1 (generated by H.D. Lipschitz) and *–*-Tubulin 4A1 (generated by M.T. Fuller) monoclonal antibodies were obtained from the Developmental Studies Hybridoma Bank, created by the NICHD of the NIH and maintained at The University of Iowa, Department of Biology, Iowa City, IA 52242.

## Extended Data Figures

**Fig. S1.**
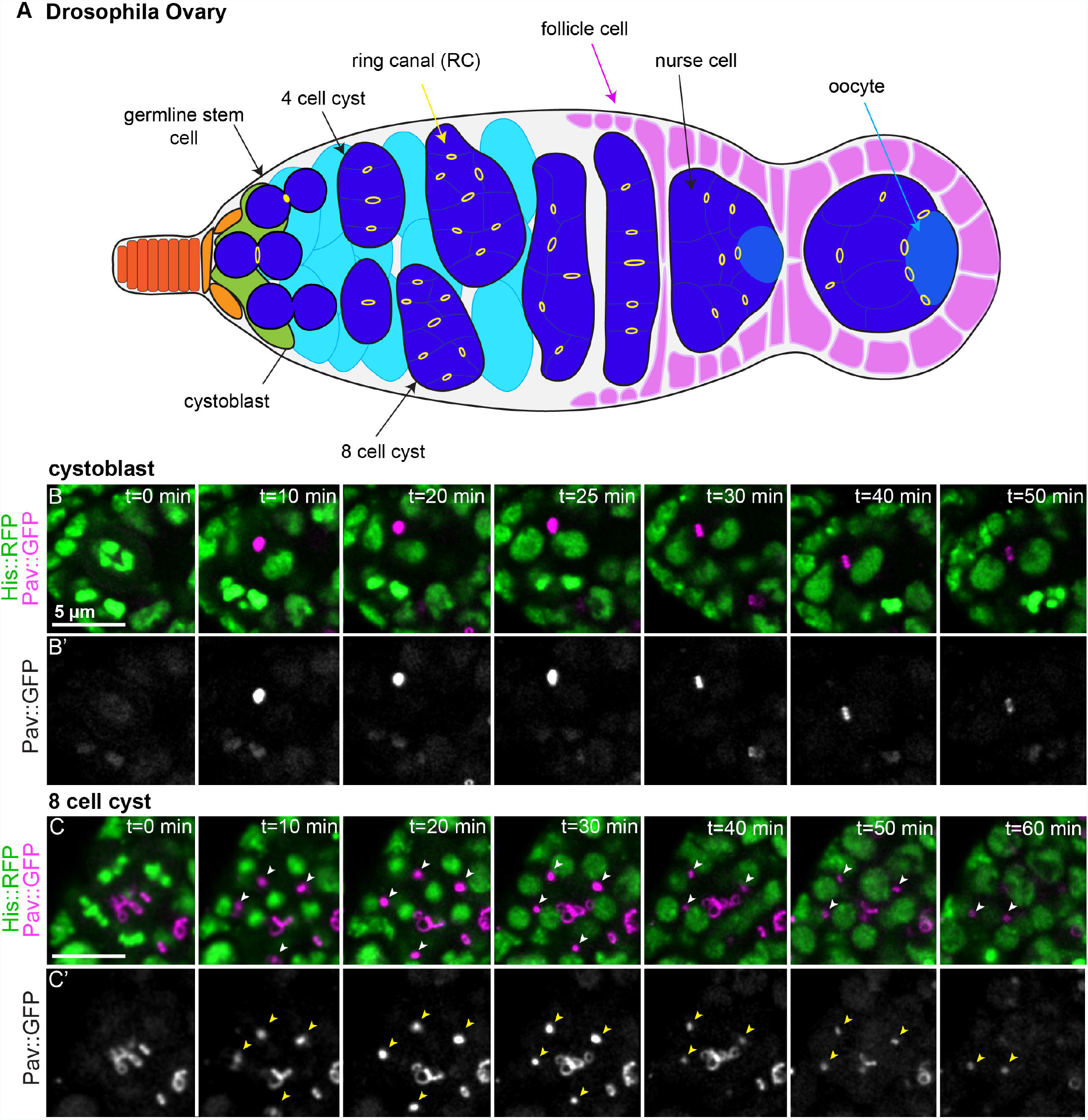
Ring canal formation in the Drosophila ovary occurs via a midbody-like intermediate. (A) Schematic of the Drosophila germarium, where four incomplete mitotic cell divisions result in a 16-cell cyst connected by 15 ring canals. Terminal filament (orange), cap cells (light orange), escort stem cells (green), escort cells (turquoise), germline cells (blue), ring canals (yellow), follicle cells (magenta), oocyte (light blue). The cystoblast divides to produce a two-cell germline cyst that divides three times more resulting in a 16-cell cyst. (B) Live imaging of a cystoblast division; a ring canal with open lumen is observed at t=50 minutes. (C) A dividing 8-cell cyst, four dividing cells are in the plane of focus as marked by four metaphase plates (t=0 minutes). The midbodies that result are marked with arrowheads and tracked over the course of the movie; at t=40 minutes, one midbody has moved out of the plane of focus, and by t=60 minutes two ring canals are in focus.

**Fig. S2.**
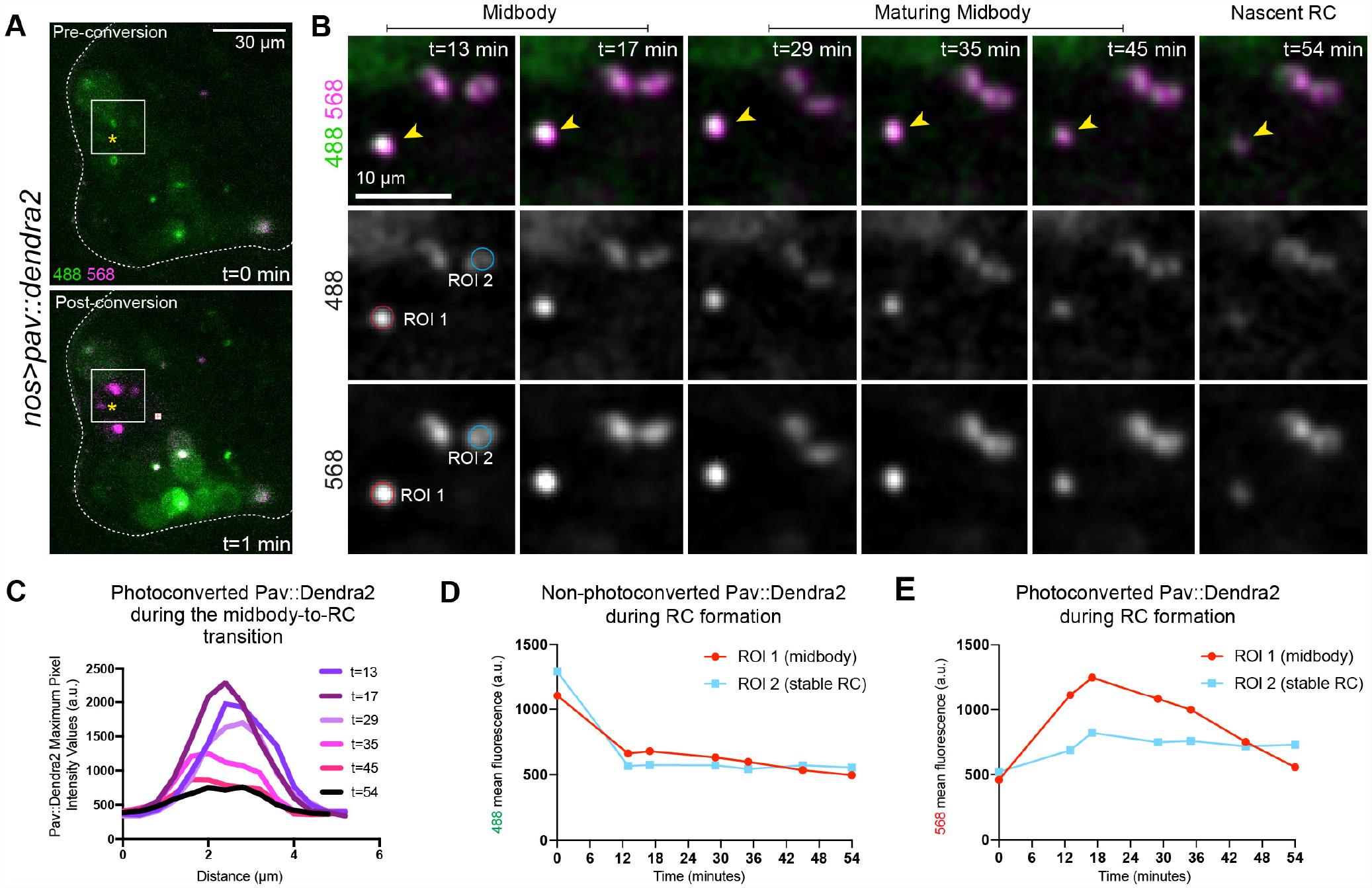
Imaging of Pav::Dendra2 reveals that ring canals are derived from germline midbodies. (A) *nos>pav::dendra2* testes before photoconversion (top, green) and after photoconversion (bottom, magenta). The photoconverted region of interest is outlined in the white box; the yellow asterisk marks the cleavage furrow of a dividing spermatogonial cell. Photoconversion occuried during cleavage furrow formation, prior to accumulation of Pav::Dendra2 in the midbody. (B) Stills from a time-lapse movie of the testis in (A) following photoconversion. (C) Fluorescence intensity across the center of Pav::Dendra2 in the midbody and ring canal of the representative sample in (A-B) over the course of time-lapse imaging. Photoconverted Pav::Dendra2 undergoes the same changes in fluorescence intensity as endogenous Pav and a ring canal localization of Pav::Dendra2 is marked at t=54 minutes. (D) Mean fluorescence intensities of a representative non-photoconverted midbody (ROI 1) and ring canal (ROI 2) imaged with 488 nm wavelength demonstrates that photoconversion of GFP-to-RFP is concomitant with a decrease in GFP signal, as expected. Following conversion, the fluorescence of GFP remains mostly stable demonstrating that no new Pav protein is added to the midbody or ring canal over the course of imaging. (E) Mean fluorescence intensities of the same representative midbody (ROI 1) and ring canal (ROI 2) reveals that the levels of photoconverted Pav::Dendra2 increase upon photoconversion and formation of the midbody and then decrease during ring canal formation whereas the levels of photoconverted Pav::Dendra2 in the ring canal are unchanged over the course of imaging.

**Fig. S3.**
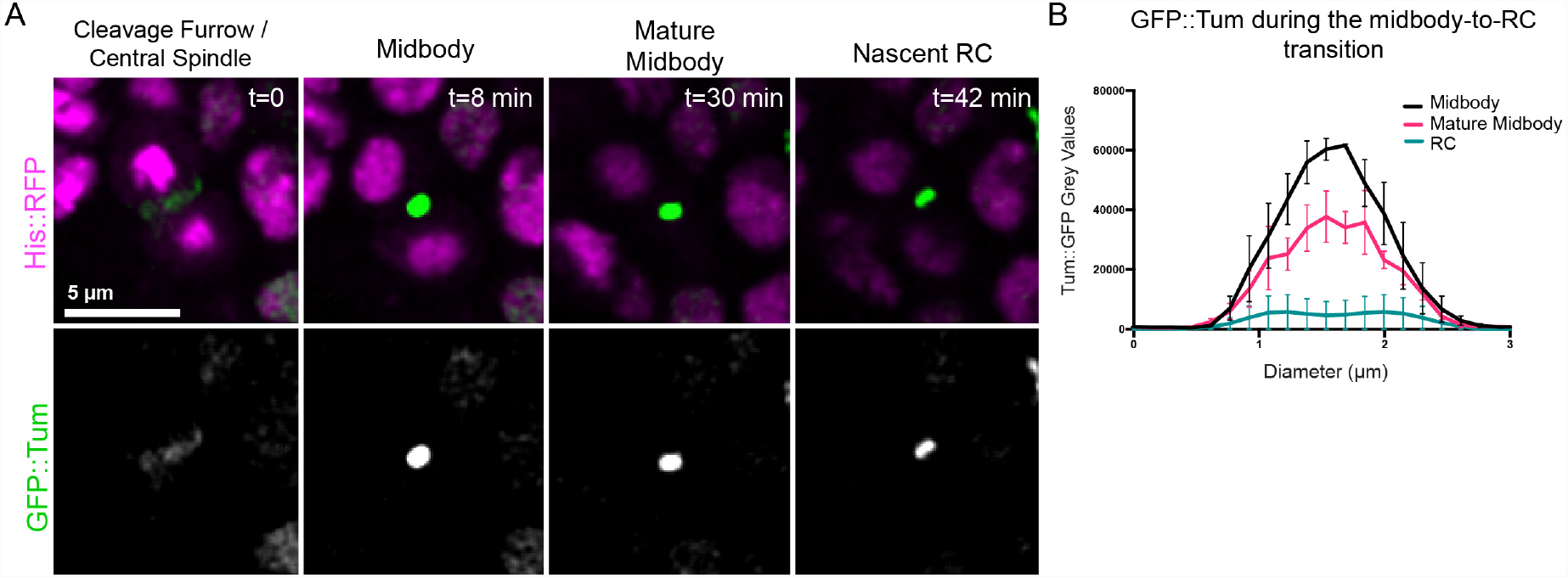
GFP::Tum dynamics during ring canal formation. (A) Single Z-slices from a time-lapse movie of a *Drosophila* testis expressing His::RFP and GFP::Tum. (B) GFP::Tum fluorescence at the indicated timepoints reveals that GFP::Tum undergoes a period of midbody maturation where the fluorescence signal is reduced *≥*5 fold, similar to Pav::mCherry.

**Fig. S4.**
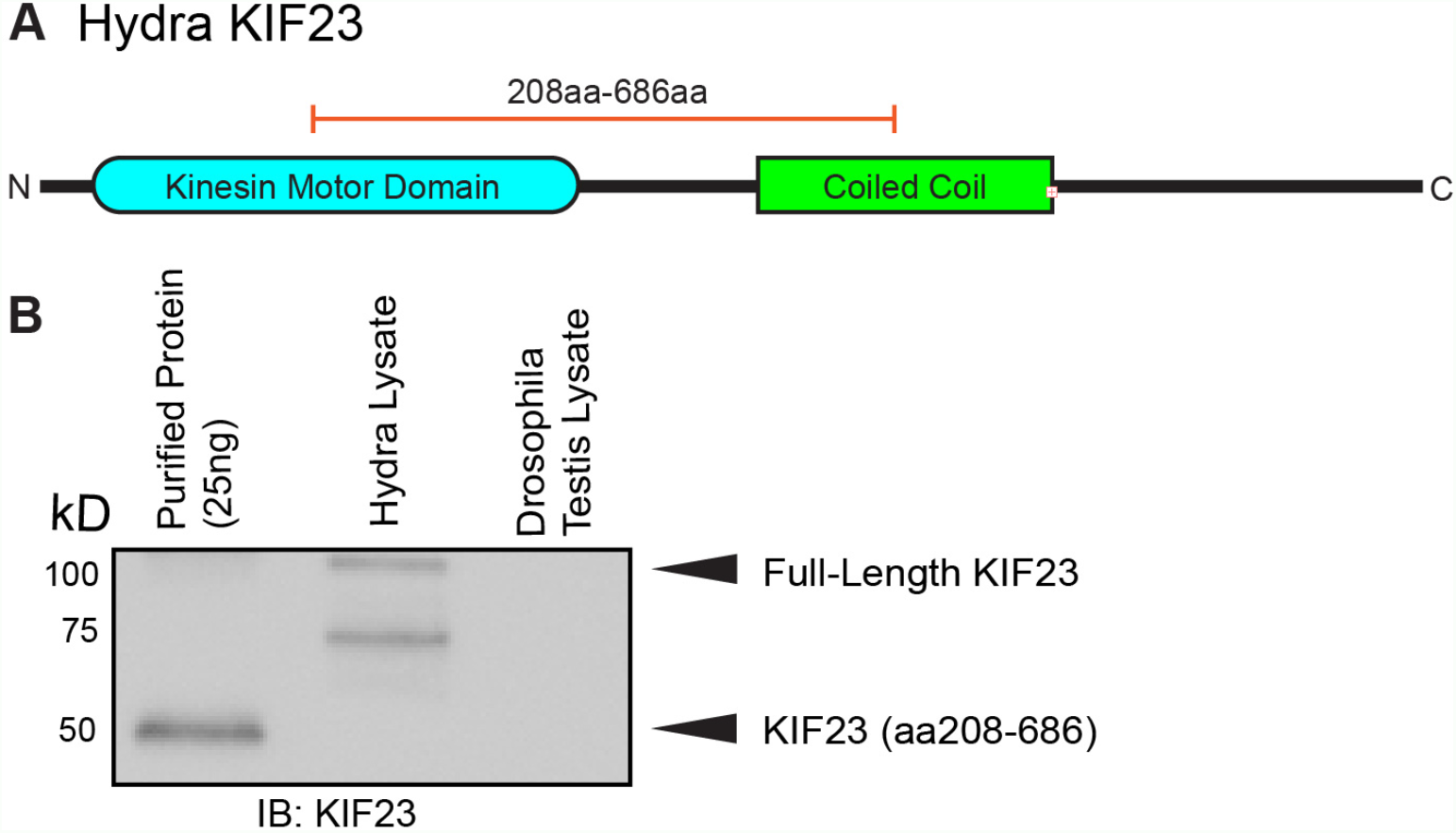
Validation of *Hydra* KIF23 polyclonal antibody. (A) Schematic of *Hydra* KIF23 protein (105 kD, predicted). Amino acids 208-686 (orange bar) were used as immunogen to generate a polyclonal antibody. This strategy was similar to the strategy used to make an antibody to *Drosophila* Pav ((20)). (B) Western blot of the purified fragment (208-686aa), *Hydra* whole cell lysate, and *Drosophila* testis whole cell lysate probed with *Hydra* anti-KIF23 antibody.

**Fig. S5.**
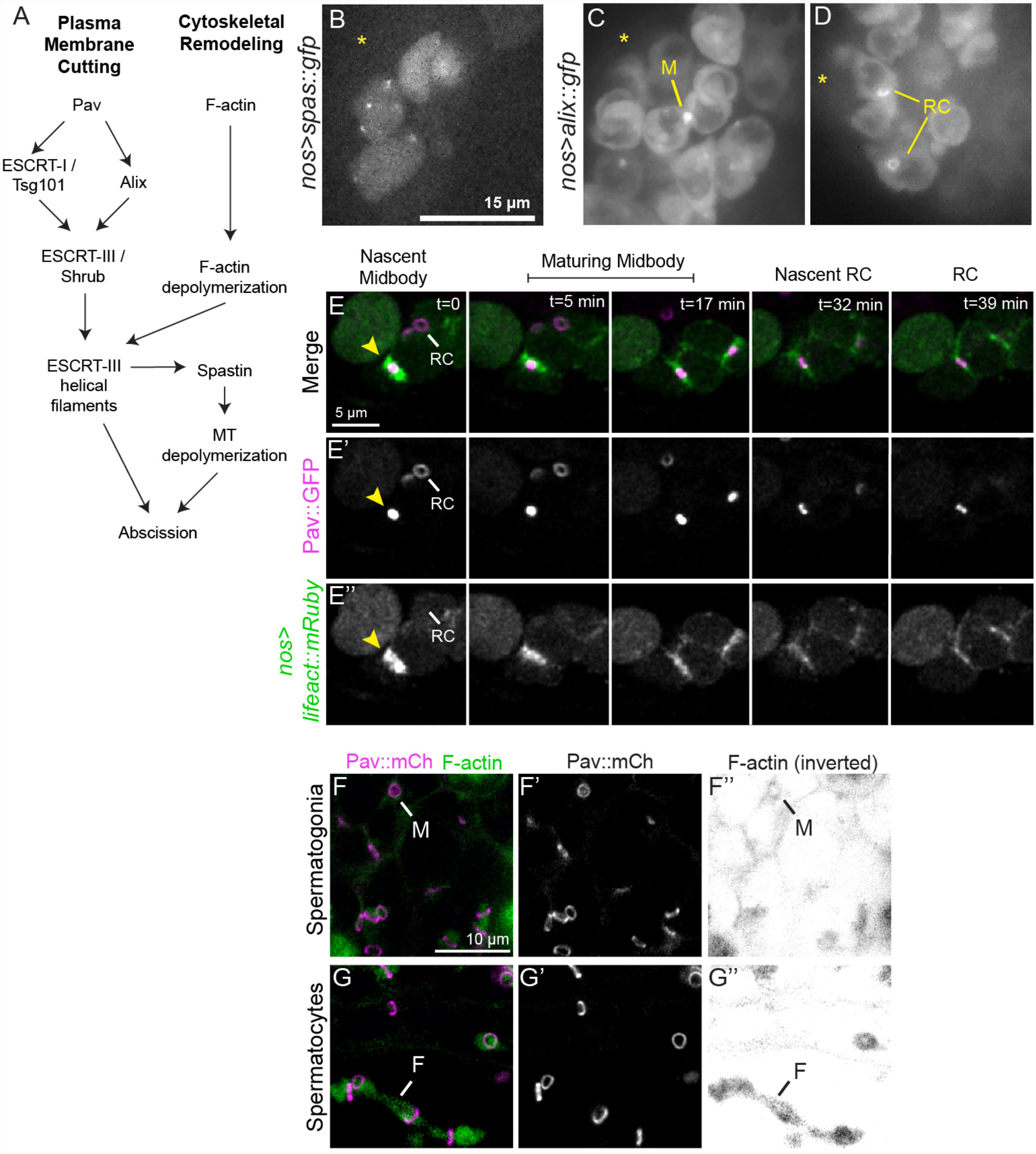
Abscission inhibition occurs at the germline midbody. (A) A pathway of abscission in *Drosophila* based on studies of complete cytokinesis and modeled after Petsalaki and Zachos (2019). (B) *nos>spas::gfp* testis; the hub is marked with an asterisk. The microtubule severase Spastin::GFP does not localize to a specific cellular structure, but as small foci. (C, D) *nos>alix::gfp* testis; the hub is marked with an asterisk. The ESCRT-associated protein Alix::GFP localizes to a midbody focus (“M”) and transiently to ring canals (“RC”) when over-expressed. (E-E”) Time-lapse imaging of *nos>lifeact::mRuby* and Pav::GFP during ring canal formation. LifeAct::mRuby is detected in the contractile ring (yellow arrowhead; t=0 minutes) and is progressively disassembled during ring canal formation such that the nascent ring canal has reduced levels of LifeAct::mRuby (t=39 minutes) and mature ring canals (ring canal (“RC”) marked in the t=0 minutes frame) have no detectable LifeAct::mRuby signal. (F-G”) Spermatogonial cells expressing Pav::mCherry were fixed and labeled with phalloidin (green) to visualize F-actin. (F-F”) At the midbody (“M”) stage, a ring of F-actin surrounds the Pav::mCherry focus, but is absent from open ring canals. (G) In spermatocytes, F-actin is detected in the fusome (“F”) but not ring canals.

## References

1. D Barthel, A Detmer, The spermatogenesis of Halichondria panicea (Porifera, Demospongiae). Zoomorphology 110, 9–15 (1990).

2. S Bertho, et al., Zebrafish dazl regulates cystogenesis and germline stem cell specification during the primordial germ cell to germline stem cell transition. Dev. (Cambridge) 148 (2021).

3. M Dym, DW Fawcett, Further observations on the numbers of spermatogonia, spermatocytes, and spermatids connected by intercellular bridges in the mammalian testis. Biol. reproduction 4, 195–215 (1971).

4. DW Fawcett, S Ito, D Slautterback, The Occurrence of Intercellular Bridges in Groups of Cells Exhibiting Synchronous Differentiation. The J. Biophys. Biochem. Cytol. 5, 453–460 (1959).

5. MP Greenbaum, T Iwamori, GM Buchold, MM Matzuk, Germ cell intercellular bridges. Cold Spring Harb. Perspectives Biol. 3, 1–18 (2011).

6. K Haglund, IP Nezis, H Stenmark, Structure and functions of stable intercellular bridges formed by incomplete cytokinesis during development (2011).

7. K Lu, L Jensen, L Lei, YM Yamashita, Stay Connected: A Germ Cell Strategy (2017).

8. D Robinson, L Cooley, Robinson and Cooley - 1996 - Stable intercellular bridges in development: the cytoskeleton lining the tunnel. Trends Cell Biol. 6, 474–479 (1996).

9. JM Mullins, JJ Biesele, Terminal phase of cytokinesis in D 98S cells. J. Cell Biol. 73, 672–684 (1977).

10. M Agromayor, J Martin-Serrano, Knowing when to cut and run: Mechanisms that control cytokinetic abscission. Trends Cell Biol. 23, 433–441 (2013).

11. JP Fededa, DW Gerlich, Molecular control of animal cell cytokinesis. Nat. Cell Biol. 14, 440– 447 (2012).

12. RA Green, E Paluch, K Oegema, Cytokinesis in animal cells. Annu. Rev. Cell Dev. Biol. 28, 29–58 (2012).

13. EA Koch, RC King, Further studies on the ring canal system of the ovarian cystocytes of Drosophila melanogaster. Zeitschrift für Zellforschung und Mikroskopische Anat. 102, 129– 152 (1969).

14. AP Mahowald, The formation of ring canals by cell furrows in Drosophila. Zeitschrift für Zellforschung und Mikroskopische Anat. 118, 162–167 (1971).

15. T Iwamori, et al., TEX14 Interacts with CEP55 To Block Cell Abscission (2010).

16. MP Greenbaum, L Ma, MM Matzuk, Conversion of midbodies into germ cell intercellular bridges. Dev. Biol. 305, 389–396 (2007).

17. GR Hime, JA Brill, MT Fuller, Assembly of ring canals in the male germ line from structural components of the contractile ring. J. Cell Sci. 109, 2779–2788 (1996).

18. SK Ong, C Foote, C Tan, Mutations of DMYPT cause over constriction of contractile rings and ring canals during Drosophila germline cyst formation. Dev. Biol. 346, 161–169 (2010).

19. YM Yamashita, Subcellular specialization and organelle behavior in germ cells (2018).

20. RR Adams, AA Tavares, A Salzberg, HJ Bellen, DM Glover, pavarotti encodes a kinesin-like protein required to organize the central spindle and contractile ring for cytokinesis. Genes Dev. 12, 1483–1494 (1998).

21. RS Kaufman, et al., Drosophila sperm development and intercellular cytoplasm sharing through ring canals do not require an intact fusome. Dev. (Cambridge) 147 (2020).

22. SK Ong, C Tan, Germline cyst formation and incomplete cytokinesis during Drosophila melanogaster oogenesis. Dev. Biol. 337, 84–98 (2010).

23. S Kuznetsov, M Lyanguzowa, TC Bosch, Role of epithelial cells and programmed ceil death in Hydra spermatogenesis. Zoology 104, 25–31 (2001).

24. S Siebert, et al., Stem cell differentiation trajectories in Hydra resolved at single-cell resolution. Science 365 (2019).

25. CK Hu, M Coughlin, TJ Mitchison, Midbody assembly and its regulation during cytokinesis. Mol. Biol. Cell 23, 1024–1034 (2012).

26. EP Karasmanis, et al., A Septin Double Ring Controls the Spatiotemporal Organization of the ESCRT Machinery in Cytokinetic Abscission. Curr. Biol. 29, 2174–2182.e7 (2019).

27. ZI Bassi, M Audusseau, MG Riparbelli, G Callaini, PP D’Avino, Citron kinase controls a molecular network required for midbody formation in cytokinesis. Proc. Natl. Acad. Sci. United States Am. 110, 9782–9787 (2013).

28. PP D’Avino, Citron kinase - renaissance of a neglected mitotic kinase. J. Cell Sci. 130, 1701– 1708 (2017).

29. PP D’Avino, L Capalbo, Regulation of midbody formation and function by mitotic kinases. Semin. Cell Dev. Biol. 53, 57–63 (2016).

30. F DiCunto, et al., Essential role of citron kinase in cytokinesis of spermatogenic precursors. J. Cell Sci. 115, 4819–4826 (2002).

31. V Naim, S Imarisio, F Di Cunto, M Gatti, S Bonaccorsi, Drosophila citron kinase is required for the final steps of cytokinesis. Mol. Biol. Cell 15, 5053–5063 (2004).

32. M Buszczak, et al., The carnegie protein trap library: A versatile tool for drosophila developmental studies. Genetics 175, 1505–1531 (2007).

33. N Karpova, Y Bobinnec, S Fouix, P Huitorel, A Debec, Jupiter, a newDrosophila protein associated with microtubules. Cell Motil. Cytoskelet. 63, 301–312 (2006).

34. M Glotzer, The 3Ms of central spindle assembly: Microtubules, motors and MAPs. Nat. Rev. Mol. Cell Biol. 10, 9–20 (2009).

35. K Murthy, P Wadsworth, Dual role for microtubules in regulating cortical contractility during cytokinesis. J. Cell Sci. 121, 2350–2359 (2008).

36. C Florindo, et al., Human Mob1 proteins are required for cytokinesis by: Controlling microtubule stability. J. Cell Sci. 125, 3085–3090 (2012).

37. JG Carlton, J Martin-Serrano, Parallels between cytokinesis and retroviral budding: A role for the ESCRT machinery. Science 316, 1908–1912 (2007).

38. E Morita, et al., Human ESCRT and ALIX proteins interact with proteins of the midbody and function in cytokinesis. EMBO J. 26, 4215–4227 (2007).

39. Wm Zhao, A Seki, G Fang, Cep55, a Microtubule-bundling Protein, Associates with Centralspindlin to Control the Midbody Integrity and Cell Abscission during Cytokinesis. Mol. biology cell 17, 3881–3896 (2006).

40. D Yang, et al., Structural basis for midbody targeting of spastin by the ESCRT-III protein CHMP1B. Nat. Struct. Mol. Biol. 15, 1278–1286 (2008).

41. A Lie-Jensen, et al., Centralspindlin Recruits ALIX to the Midbody during Cytokinetic Abscission in Drosophila via a Mechanism Analogous to Virus Budding. Curr. Biol. 29, 3538– 3548.e7 (2019).

42. ÅH Eikenes, et al., ALIX and ESCRT-III Coordinately Control Cytokinetic Abscission during Germline Stem Cell Division In Vivo. PLoS Genet. 11 (2015).

43. E Petsalaki, G Zachos, Building bridges between chromosomes: novel insights into the abscission checkpoint (2019).

44. SJ Terry, F Donà, P Osenberg, JG Carlton, US Eggert, Capping protein regulates actin dynamics during cytokinetic midbody maturation. Proc. Natl. Acad. Sci. United States Am. 115, 2138–2143 (2018).

45. ÅH Eikenes, A Brech, H Stenmark, K Haglund, Spatiotemporal control of Cindr at ring canals during incomplete cytokinesis in the Drosophila male germline. Dev. Biol. 377, 9–20 (2013).

46. DN Robinson, K Cant, L Cooley, Morphogenesis of Drosophila ovarian ring canals. Development 120, 2015–2025 (1994).

47. EF Oakberg, Duration of spermatogenesis in the mouse and timing of stages of the cycle of the seminiferous epithelium. Am. J. Anat. 99, 507–16 (1956).

48. MT Fuller, The Development of Drosophila melanogaster eds. M Bate, A Martinez Arias. (Cold Spring Harbor Laboratory Press, Plainview, New York), 1 edition, pp. 71–148 (1993).

49. YH Inoue, et al., Mutations in orbit/mast reveal that the central spindle is comprised of two microtubule populations, those that initiate cleavage and those that propagate furrow ingression. J. Cell Biol. 166, 49–60 (2004).

50. N El-Amine, A Kechad, S Jananji, GR Hickson, Opposing actions of septins and Sticky on Anillin promote the transition from contractile to midbody ring. J. Cell Biol. 203, 487–504 (2013).

51. FT Bianchi, et al., Citron Kinase Deficiency Leads to Chromosomal Instability and TP53Sensitive Microcephaly. Cell Reports 18, 1674–1686 (2017).

52. FT Bianchi, M Gai, GE Berto, F Di Cunto, Of rings and spines: The multiple facets of Citron proteins in neural development. Small GTPases 11, 122–130 (2020).

53. H Li, et al., Biallelic Mutations in Citron Kinase Link Mitotic Cytokinesis to Human Primary Microcephaly. Am. J. Hum. Genet. 99, 501–510 (2016).

54. P Madaule, et al., Role of citron kinase as a target of the small GTPase Rho in cytokinesis. Nature 394, 491–494 (1998).

55. KC McNeely, ND Dwyer, Cytokinetic Abscission Regulation in Neural Stem Cells and Tissue Development. Curr. Stem Cell Reports (2021).

56. MJ Dayel, et al., Cell differentiation and morphogenesis in the colony-forming choanoflagellate Salpingoeca rosetta. Dev. Biol. 357, 73–82 (2011).

57. H Hoops, et al., Cytoplasmic Bridges in Volvox and Its Relatives in Cell-Cell Channels. pp. 65–84 (2006).

58. KF Lenhart, S DiNardo, Somatic Cell Encystment Promotes Abscission in Germline Stem Cells following a Regulated Block in Cytokinesis. Dev. Cell 34, 192–205 (2015).

59. LX Morris, AC Spradling, Long-term live imaging provides new insight into stem cell regulation and germline-soma coordination in the Drosophila ovary. Development 138, 2207–2215 (2011).

60. P. D’Avino, MS Savoian, DM Glover, Mutations in sticky lead to defective organization of the contractile ring during cytokinesis and are enhanced by Rho and suppressed by Rac. J. Cell Biol. 166, 61–71 (2004).

